# Development of circadian immune regulation in early life

**DOI:** 10.64898/2026.06.04.730046

**Authors:** Xia Li, Andrew Kwon, Zeynep Melis Gül, Paula Rothämel, Florian Seytter, Lou Martha Wackerbarth, Sarah Kim-Hellmuth, Claudia Nussbaum, Shaon Sengupta, Markus Sperandio, Christoph Scheiermann

**Affiliations:** Institute of Cardiovascular Physiology and Pathophysiology, Biomedical Center, Ludwig-Maximilians-Universität München, Planegg-Martinsried, Germany; Department of Neuroscience, University of Pennsylvania, University of Pennsylvania, Philadelphia, USA; Department of Pathology and Immunology, Faculty of Medicine, University of Geneva, Geneva, Switzerland; Division of Neonatology, Department of Pediatrics, Dr. von Hauner Children’s Hospital, LMU Medizin, LMU University Hospital Ludwig-Maximilians-University Munich; Computational Health Center, Institute of Translational Genomics, Helmholtz Munich, Neuherberg, Germany; University of Pennsylvania Perelman School of Medicine, Philadelphia, USA; The Children’s Hospital of Philadelphia, Philadelphia, USA; Translational Research Centre in Onco-Hematology (CRTOH), Geneva, Switzerland; Geneva Centre for Inflammation Research (GCIR), Geneva, Switzerland; Institute of Genetics and Genomics of Geneva (iGE3), Geneva, Switzerland

## Abstract

Circadian rhythms temporally regulate immune cell trafficking, abundance, and immune responsiveness in adults, yet how this rhythmic organization emerges during postnatal development remains unknown. Here, we systematically investigated the establishment and development of circadian immune regulation across multiple biological levels. Leukocyte rhythmicity was not present at birth but was gradually established and strengthened during postnatal development in a leukocyte subset-dependent manner. In parallel, promigratory markers displayed dynamic and heterogeneous developmental trajectories, indicating progressive coordination of rhythmic promigratory programs. At the systemic level, peripheral clocks matured and synchronized during development, with tissue- and clock-gene-dependent differences in the acquisition timing, peak phase, and amplitude of circadian oscillations. This multi-level analysis identifies postnatal development as a dynamic period during which circadian immunity is progressively consolidated through leukocyte rhythmicity, regulation of promigratory factors, and peripheral clocks.

## Introduction

Circadian rhythms align physiological processes with the 24 h light-dark environment and have emerged as important regulators of immune homeostasis and responses^1,2^. In adults, both leukocyte abundance and trafficking behavior exhibit pronounced time-of-day-dependent rhythmicity^3^. These rhythms are driven by both cell-intrinsic molecular clocks and rhythmic promigratory cues derived from immune cells, peripheral tissues, and the environment. At the molecular level, transcription-translation feedback loops formed by core clock genes/proteins generate intrinsic circadian oscillations within immune cells, whereas migration- and adhesion-related factors contribute to rhythmic leukocyte circulation and migration at the organismal level^4,5^. Together with systemic timing cues, these mechanisms form the basis of circadian immune rhythmicity in adulthood. By incorporating circadian timing into immune interventions, temporal regulation can strongly influence inflammatory responses and the efficacy of therapeutic strategies^6,7^.

In contrast to the adult immune system, neonatal immunity displays distinct characteristics under both steady state and inflammatory conditions^8-10^. Immediately after birth, the immune system adapts from a relatively sterile environment to an antigen-rich external world, with innate immune parameters dominating the earliest phase of postnatal life^11,12^. This is followed by extensive remodeling of immune composition, tissue physiology, metabolism, and environmental entrainment, a period likely associated with the emergence of rhythmic immune regulation^13^. However, it remains unclear how circadian immune organization develops during this critical window. In particular, it is unknown whether circadian immune rhythms are already established at birth or are gradually assembled during postnatal development. To address these questions, we investigated postnatal circadian immune development in mice by profiling leukocyte abundance, promigratory marker expression, and clock gene rhythms in leukocytes and peripheral tissues across the 24 h cycle at different postnatal ages. Together, this study examines postnatal development as a dynamic window for the establishment of circadian immune organization across multiple biological layers.

## Results

### Leukocyte rhythmicity is progressively established after birth

To determine when and how circadian leukocyte rhythms are established after birth, we profiled circulating leukocytes and their subsets in mice, including B cells, CD4^+^ T cells, CD8^+^ T cells, neutrophils, classical monocytes (cMono), non-classical monocytes (ncMono), NK cells and eosinophils, at multiple postnatal ages and zeitgeber time (ZT) points (ZT1, ZT7, ZT13, ZT19; hours after light onset under a 12 h light:12 h dark cycle) using hematological analyses and flow cytometry. Total leukocyte counts displayed limited oscillations immediately after birth (**Fig. 1A**). During postnatal development, overall leukocyte counts increased and exhibited more robust daily rhythmicity at later ages around postnatal day (P) 17, indicating that circadian regulation of circulating leukocyte abundance is developmentally acquired rather than fully established at birth. The selected leukocyte subsets showed similar subset-specific developmental changes in daily rhythmicity. Neutrophils (**Fig. 1B**) and cMono (**Fig. 1E**) were already abundant during early postnatal life, whereas B cells (**Fig. 1C**) and CD4^+^ T cells (**Fig. 1D**) became more prominent later, with innate immune cells being reduced at the same time. However, none of these subsets showed a clearly established daily rhythm at birth. Across the examined populations, oscillatory patterns became more evident during postnatal development, with stronger time-of-day differences first observed around P17. This was followed by a transient reduction or loss of detectable rhythmicity around P21, before rhythmic patterns re-emerged at later ages. Despite this temporary weakening, peak abundance remained broadly aligned with the light phase, corresponding to the resting phase in mice. Consistent with these observations, acrophase and amplitude mapping across leukocyte subsets showed increasing oscillation strength, and progressive synchronization, which peaked preferentially during the behavioral rest phase (**Fig. 1F**). These findings demonstrate that circadian leukocyte rhythms are gradually acquired after birth and share a common developmental trajectory across major circulating leukocyte populations, with increasing oscillation strength and temporal organization.

**Figure 1.**
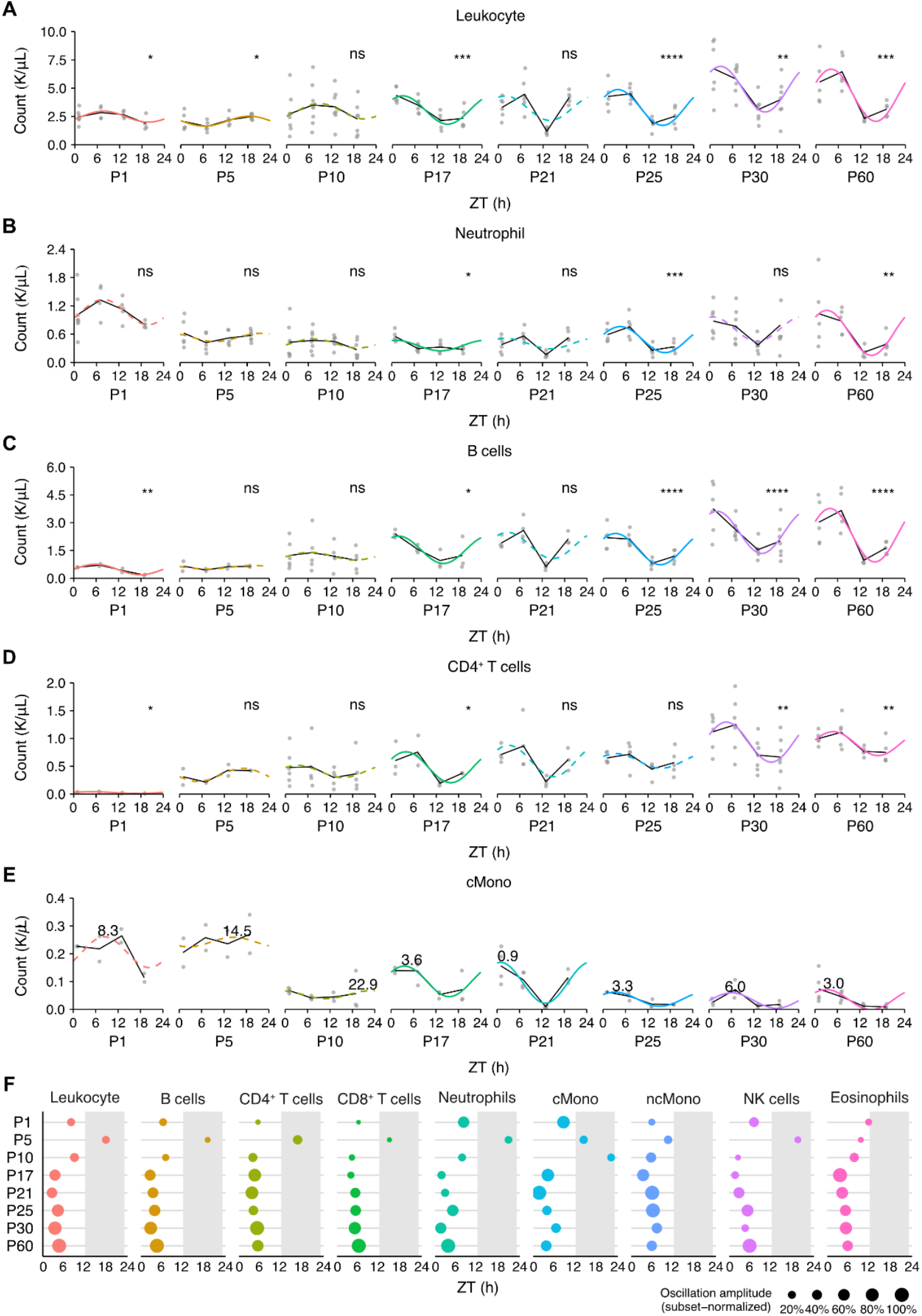
Postnatal leukocyte rhythms emerge progressively after birth and strengthen with age. Developmental profiles of circulating leukocyte counts across the day at different postnatal ages. (A) total leukocytes, n = 4–8 mice per age and ZT, (B) neutrophils, n = 4–8 mice per age and ZT, (C) B cells, n = 2–7 mice per age and ZT, (D) CD4^+^ T cells, n = 2–7 mice per age and ZT, (E) classical monocytes, n = 2–7 mice per age and ZT. For (A-E) data were analyzed using cosinor analysis. Solid lines indicate significant rhythmicity, while dashed lines indicate non-significant fitted rhythms. Significance is indicated as ns, not significant; \**p* < 0.05; \*\**p* < 0.01; and \*\*\*\**p* < 0.0001. (F) Acrophase and amplitude mapping of rhythmic leukocyte populations across postnatal ages.

### Oscillations in promigratory markers develop dynamically across postnatal stages

Circulating leukocyte oscillations in the adult immune system are closely associated with coordinated expression of migration- and adhesion-related factors^3^. Therefore, we next investigated how rhythms in the expression of promigratory markers are established and organized during postnatal development. Using flow cytometry, we quantified the daily expression pattern of CXCR4, which functions as a key regulator of leukocyte recruitment to peripheral tissues, across all leukocytes at multiple postnatal ages. The developmental profile showed a clear age-dependent progression in the rhythmic expression of CXCR4 (**Fig. 2A**). CXCR4 expression displayed weak oscillatory patterns at early postnatal ages, whereas stronger daily rhythms emerged after weaning and persisted into adulthood. We further analyzed the developmental organization of additional critical promigratory factors for leukocyte trafficking, including PSGL-1, CD62L, CD11a, CD11b, CD18, CD49d, and CXCR2, across different leukocyte subsets (**Fig. 2B**). This analysis revealed marked heterogeneity in rhythmic marker expression across these adhesion-relevant molecules for the various cell types, and postnatal ages. Although no uniform developmental pattern was observed across all markers, the amplitude of the oscillations generally increased at later ages, and peak timing became more organized in several markers. To summarize the coordinated behavior of migration- and adhesion-relevant markers, we calculated an integrated promigratory index, based on their expression profiles and analyzed its developmental pattern within each leukocyte subset. The result showed that promigratory marker oscillations became more organized during development in several leukocyte populations (**Fig. 2C**). At later ages, peak phases of the promigratory factors were frequently positioned around the light–dark transition, coinciding with the trough phase of circulating leukocyte abundance. This temporal relationship suggests that increased expression of migration- and adhesion-related programs may be associated with reduced circulating leukocyte numbers, consistent with enhanced tissue recruitment at this phase.

**Figure 2.**
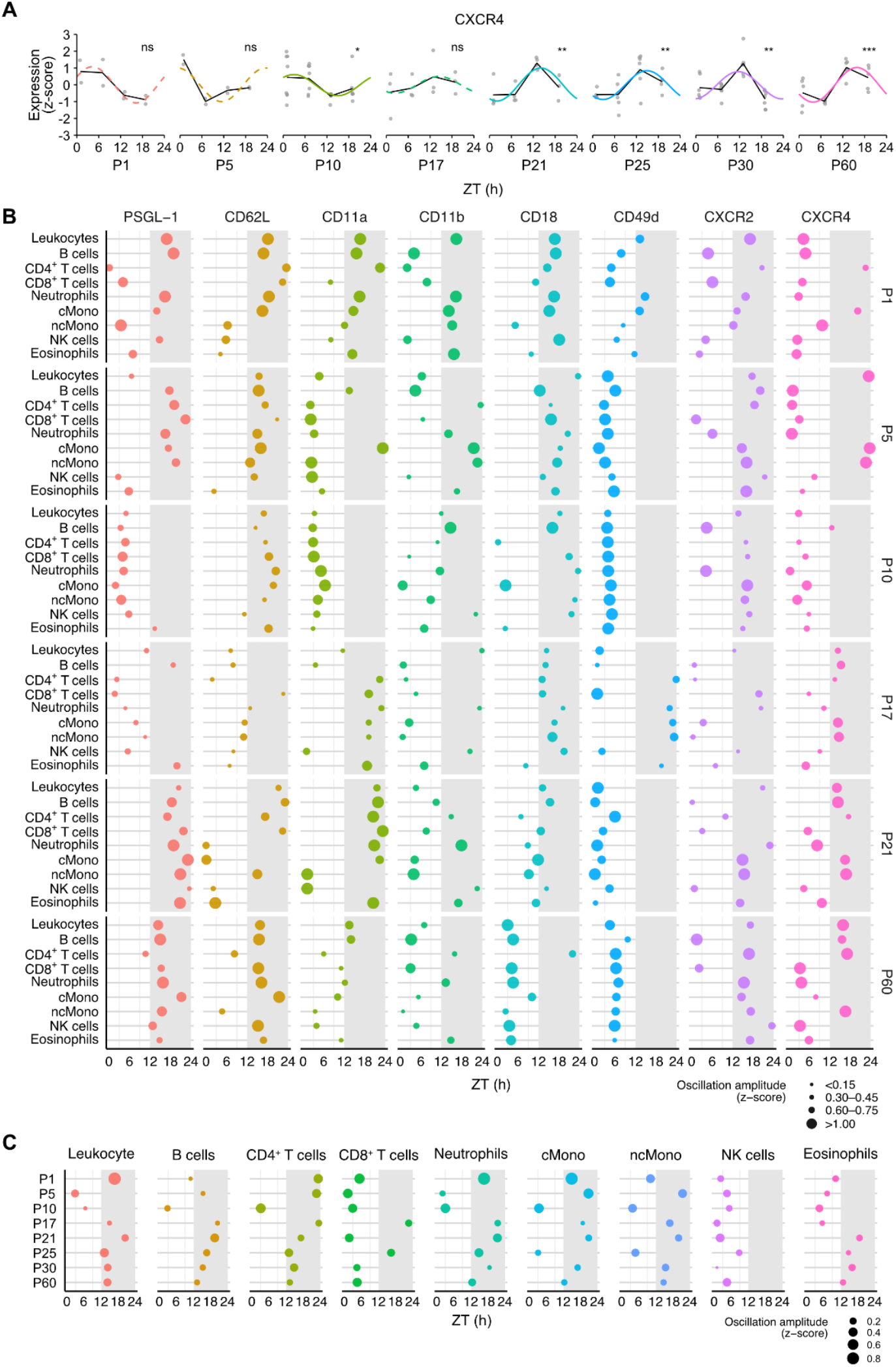
Promigratory factors show dynamic oscillatory patterns during postnatal development. (A) Daily expression profiles of CXCR4 on total circulating leukocytes at different postnatal ages. n = 2–7 mice per age and ZT. Data were processed by z-score normalization and analyzed using cosinor analysis. Solid lines indicate significant rhythmicity, while dashed lines indicate non-significant fitted rhythms. Significance is indicated as ns, not significant; \**p* < 0.05; \*\**p* < 0.01; and \*\*\*\**p* < 0.0001. (B) Acrophase and amplitude mapping of promigratory marker rhythms across leukocyte subsets and postnatal ages. (C) Acrophase and amplitude mapping of the promigratory index across leukocyte subsets and postnatal ages.

### Peripheral clocks mature and synchronize in a tissue- and gene-dependent manner

Upstream to leukocyte oscillations and promigratory marker rhythms, expression patterns of peripheral clocks represent the most important level of circadian organization. To determine whether the postnatal maturation of leukocyte rhythms was associated with the development of systemic circadian clock expression across tissues, we analyzed clock gene expression in circulating leukocytes and peripheral tissues. In leukocytes, expression of the core clock gene *Bmal1* showed apparent rhythmicity as early as P5, further developed dynamically during the postnatal period, and eventually exhibited stable oscillations with peak phases around the light-dark transition. (**Fig. 3A**). To obtain an overview of systemic rhythms, we quantified peak phase and oscillation strength of the core clock genes, including *Bmal1, Clock, Per1, Per2, Cry1, Cry2, Nr1d1, Nr1d2*, and *Dbp*, across blood leukocytes, bone marrow, spleen, brain, kidney, small intestine, liver, and heart. The result revealed widespread but heterogeneous circadian organization across tissues and genes during postnatal development (**Fig. 3B**). At an early age, rhythmic clock gene expression was relatively weak and showed variable peak timing. During development, oscillation amplitude increased for several genes and tissues, and peak phases became more consistently organized. However, this maturation was not uniform, with different organs and clock genes displaying distinct developmental trajectories. To further summarize the development of systemic clock organization, we integrated clock gene expression patterns across all analyzed tissues for each individual gene. This analysis showed that several clock genes acquired stronger and more temporally synchronized oscillatory patterns from P10, reflected by increased amplitude and more coherent systemic acrophase distributions (**Fig. 3C**). *Bmal1* and *Clock* became preferentially aligned around the transition from the dark to the light phase, whereas *Per, Cry, Nr1d*, and *Dbp* showed peak phases mainly around the light– dark transition, representing the pattern seen in adults and indicating the emergence of gene-specific phase organization across the systemic clock network. Together, these data indicate that the emergence of leukocyte rhythmicity after birth occurs alongside the progressive organization of molecular circadian rhythms across the body. This coordinated but tissue- and gene-dependent maturation may contribute to the developmental establishment of circadian immune regulation.

**Figure 3.**
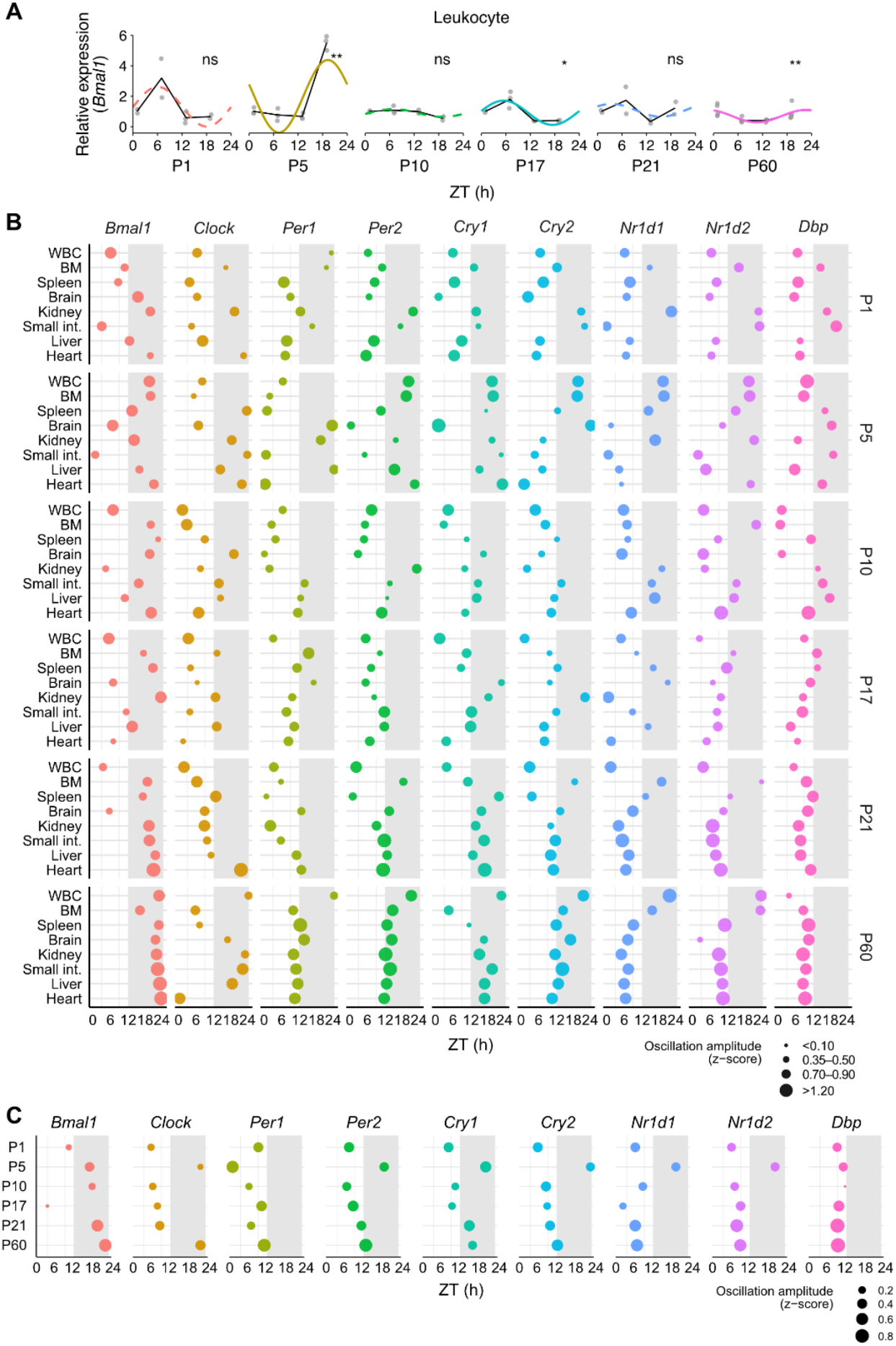
Peripheral clocks undergo progressive maturation and temporal organization during development. (A) Daily expression profiles of *Bmal1* in total circulating leukocytes across postnatal development. n = 2–5 mice per age and ZT. Data were analyzed using cosinor analysis. Solid lines indicate significant rhythmicity, while dashed lines indicate non-significant fitted rhythms. Significance is indicated as ns, not significant; \**p* < 0.05; and \*\**p* < 0.01. (B) Acrophase and amplitude mapping of clock gene expression across leukocytes, bone marrow, spleen, brain, kidney, small intestine, liver, and heart at different postnatal ages. (C) Acrophase and amplitude mapping of systemic clock gene rhythmicity across postnatal development.

## Discussion

In this study, we identify postnatal development as a period during which circadian immune organization is progressively established across multiple biological layers. Rhythmicity of circulating leukocytes was not present at birth but emerged during postnatal development across all major leukocyte populations. This process was accompanied by developmental organization of rhythmicity in migration- and adhesion-related markers and by progressive maturation of molecular clock gene expression across peripheral tissues. Together, these data indicate that the temporal organization of the circadian immune system is developmentally acquired rather than fully mature at birth. A central feature of this developmental process is heterogeneity. Rhythmicity did not emerge synchronously across leukocyte subsets, promigratory markers, tissues, or clock genes. Instead, different biological layers followed partially distinct trajectories, although the overall tendency was toward stronger and more organized oscillatory patterns with age. At the level of circulating leukocytes, the acquisition of rhythmicity followed a broadly similar pattern across the populations examined, suggesting that leukocyte rhythmicity develops as a coordinated feature of circulating immune cells rather than through the independent maturation of individual subsets. The transient weakening observed at intermediate postnatal stages, around the time of weaning, further suggests that this process is not a simple linear strengthening of oscillations.

Rhythms in migration- and adhesion-related markers add a potential mechanistic layer to the maturation of leukocyte oscillations. Because these molecules regulate leukocyte adhesion, circulation, recruitment and egress to and from tissues, their developmental rhythmicity may influence the timing of leukocyte retention in blood. The progressive organization of the promigratory index supports the idea that leukocyte rhythmicity develops together with coordinated changes in trafficking-related molecular programs.

The development of immune rhythmicity also occurred in parallel with the maturation of peripheral circadian clocks. Circulating leukocytes are exposed to rhythmic systemic and tissue-derived cues, including endocrine, metabolic, neural, and local vascular signals. Therefore, the progressive organization of clock gene expression across tissues suggests that leukocyte rhythms may depend not only on leukocyte-intrinsic timing mechanisms, but also on the developing temporal structure of the organism as a whole.

### Limitations to this study

Our analyses define developmental patterns across leukocyte counts, promigratory marker expression, and clock gene rhythms, but do not yet directly establish functional relationships among these layers. Our ongoing work will determine how leukocyte-intrinsic clocks, peripheral tissue clocks, systemic cues, and migration-related programs interact during early life. In addition, the relevance of these findings to neonatal immune activation, such as caused by vaccinations and infections, remains to be established.

Overall, our results indicate that circadian immune organization is assembled after birth through a dynamic and heterogeneous developmental process across circulating leukocytes, promigratory factors, and peripheral clocks. This should prove to be of relevance in leveraging time-of-day to optimize immune interventions in early life in infants, such as with immunizations.

## Methods

### Mice

Male and female C57BL/6NCrl mice were obtained either from Charles River Laboratories or from the Core Animal Facility of the Biomedical Center, LMU. Mice were maintained under a 12 h light:12 h dark cycle with ad libitum access to food and water. For collection at different time points within the same experimental day, mice were housed in light-controlled chambers (Park Bio) with shifted 12 h light:12 h dark cycles before sampling. To generate timed litters, male and female C57BL/6NCrl mice were set up for pair or trio mating at 8–12 weeks of age. Offspring were analyzed at four *zeitgeber* (ZT) time points (ZT1, ZT7, ZT13, and ZT19) across postnatal development (P1, P5, P10, P17, P21, P25, P30, and P60). During the dark phase, all procedures were performed under dim red light. All animal experiments were performed in accordance with the German Animal Welfare Act and were approved by the Regierung von Oberbayern.

### Sample collection

At the indicated experimental time points, blood and tissues were collected from mice across postnatal development. Mice aged P1 and P5 were euthanized by decapitation, during which blood was collected into EDTA-coated tubes. For mice aged P10 and older, blood was collected by cardiac puncture immediately after cervical dislocation. Tissues (brain, heart, liver, spleen, kidney, and small intestine) were harvested immediately after euthanasia, placed in RNAprotect Tissue Reagent (QIAGEN) at 4°C overnight, and further stored at −20°C for RNA isolation. Bone marrow was collected by flushing femurs and tibias with cold PBS and immediately processed for RNA isolation.

### Hematological analysis and flow cytometry

Blood samples were subjected to complete blood count analyses using an IDEXX Procyte DX cell counter. Afterwards, erythrocytes were lysed by red blood cell lysis buffer (0.8% NH_4_Cl) 2 times on ice, for 10 min each. The lysis was stopped by adding an equal amount of PBS. After centrifugation, cells were resuspended in PBS, and stained with Zombie UV Fixable Viability Kit (BioLegend) for 20 min at room temperature in the dark. Cells were further washed, resuspended in PEB (PBS supplemented with 2% fetal bovine serum and 2mM EDTA), and stained with fluorescence-conjugated antibodies for 40 min on ice. After staining, cells were fixed with 2% PFA for 15 min on ice, washed with PBS and resuspended in 300 μL PEB for flow cytometry analysis using Cytek Aurora (Cytek Biosciences).

### RNA isolation, reverse transcription, and qPCR

Total RNA was extracted from blood and bone marrow samples following red blood cell lysis, as well as from tissue samples, using RNAeasy Protect Mini kit (QIAGEN) or RNeasy Mini Kit (QIAGEN) according to the manufacturer’s instructions. RNA quality and quantity were measured using a NanoDrop 2000 spectrophotometer (Thermo Fisher Scientific). Reverse transcription was performed to obtain cDNA samples using High-Capacity cDNA Reverse Transcription Kit (Thermo Fisher Scientific) by following the provided instructions. Q-PCR was performed using PowerUp SYBR Green Master Mix (Thermo Fisher Scientific) by QuantStudio 6 Pro Real-Time PCR System (Thermo Fisher Scientific) or StepOnePlus Real-Time PCR System (Thermo Fisher Scientific).

### Data processing and statistical analysis

Flow cytometry data were analyzed using FlowJo. Absolute cell numbers were calculated by multiplying total WBC counts obtained from hematological analysis by the corresponding subset frequencies. For promigratory marker analyses, median fluorescence intensity (MFI) values were extracted for each marker within each leukocyte subset. Raw MFI values were corrected for batch effects by subtracting the batch-specific median value and were subsequently z-score normalized within each postnatal age, leukocyte subset, and marker. A promigratory index was calculated for each sample as the mean z-scored expression of the selected migration- and adhesion-related markers.

For clock gene expression analysis, gene expression levels were analyzed using the 2^-ΔΔCt^ method and normalized to the reference genes Rpl32 or Rplp0. For systemic clock analysis, clock gene expression values were z-score normalized within each tissue and gene before integration across all analyzed tissues for each individual gene.

Daily rhythmicity was assessed using a 24 h cosinor model with cosine and sine terms fitted to ZT. Rhythmicity was tested by comparing the full cosinor model with an intercept-only null model. Amplitude and acrophase were derived from the fitted model and expressed in relative units and ZT hours respectively.

Statistical analysis and data visualization were performed in R. All data are shown as mean. Significance is indicated as ns, not significant; \**p* < 0.05; \*\**p* < 0.01; \*\*\**p* < 0.001; \*\*\*\**p* < 0.0001.

## Data availability

Source data and analysis code will be made available upon reasonable request at the time of final publication.

## Acknowledgements

We thank the Core Flow Cytometry Facility and the Core Animal Facility of the Biomedical Center, LMU, for technical assistance and animal care. This work was supported by the German Research Foundation (DFG) collaborative research grant TRR359 (#491676693; projects B02 to M.S. and R.I., B06 to S. K-H. and C.N., and B07 to M.S. and C.S.) and TRR418 (#541063275; project A01 to C.S.)). This study was further supported by the European Research Council (ERC CoG 101001233, CIRCADYN), the Swiss National Science Foundation (SNSF 310030_219256, 10.000.652 and IZJA-3_238772 / 1 to C.S and 215050 to D.M), Swiss Cancer Research (KFS-5898-08-2023), the Geneva Cancer League (2403), the Translational Research Center in Oncohaematology (CRTOH) (2024-CRTOH-GTO_SA_24_003, a donation to the GTO programme, the Fondation Dr Henri Dubois-Ferrière Dinu Lipatti (DFDL), and the Fondation privée of the Geneva University Hospitals to C.S.) and the Medical & Clinician Scientist Program (MCSP) of the Medical Faculty of the Ludwig-Maximilians University Munich to C.N.

